# Daple-FLT3 (CCDC88C-FLT3) gene fusion requires the coiled-coil domain for maximal activation and pericentrosomal localization

**DOI:** 10.1101/2025.09.25.678166

**Authors:** Darren Kao, Michael Acquazzino, Arnel Ibarra, Elena Valenzuela, Haley Mai, Kimberly Aguilar, Elliot Stieglitz, Jason Ear

## Abstract

Gene fusions are stable protein products often occurring from chromosomal rearrangements. These chimeric proteins typically contain distinct molecular entities from each parent gene, and thus, create a product with altered or aberrant function. Gene fusions are frequently found in cancers, including Leukemia. Here, we characterize the kinase activity and subcellular distribution of the Daple-FLT3 (CCDC88C-FLT3) fusion oncoprotein—a rare, but recurrent gene fusion found in patients with hematological malignancies. The protein contains the FLT3 kinase domain and is activated without ligand stimulation. This leads to activation in STAT5a, AKT, and MAPK signaling, which can be modulated by the tyrosine kinase inhibitors (TKIs) sorafenib, quizartinib, and to a lesser degree, imatinib. Moreover, fusion of this kinase domain to Daple facilitates its localization to the pericentrosomal space and enhances kinase activation. These findings provide evidence that targeting Daple-FLT3 outside of its kinase domain may be a complementary approach with TKI therapy.

**Key Points:**

- Daple-FLT3 fusion proteins contain a constitutively active kinase domain, activating distinct signaling molecules in cells
- Coiled-coil domain on Daple is dispensable for kinase activation, but necessary for maximal activation

## Introduction

FLT3 is a receptor tyrosine kinase that regulates cellular properties such as proliferation, survival, and differentiation. Various FLT3 alterations including internal tandem duplication (ITD), point mutations, and chromosomal rearrangements have been found in leukemia and other myeloproliferative disorders^1–3^. Compared to the other alterations, chromosomal rearrangements involving FLT3 is quite rare^4–6^, but is being identified more frequently as RNA sequencing become more routine in cancer subtyping^7,8^.

Chromosomal rearrangements between Daple (CCDC88C) and FLT3 have been identified in two patients, one with Juvenile myelomonocytic leukemia (JMML) and another with a myeloid/lymphoid neoplasm (MLN)^9,10^. One patient carried a chromosomal rearrangement where Daple on Exon23 was partnered to FLT3 on Exon14. Another patient had a rearrangement between Daple on Exon19 and FLT3 on Exon14 (Figure 1A). We refer to these fusions as Daple-FLT3 Ex23-14 and Daple-FLT3 Ex19-14. In both cases, the chimeric protein contain Daple’s N-terminal HOOK and coiled-coil domain rearranged to the tyrosine kinase domain of FLT3 (Figure 1B).

**Figure 1.**
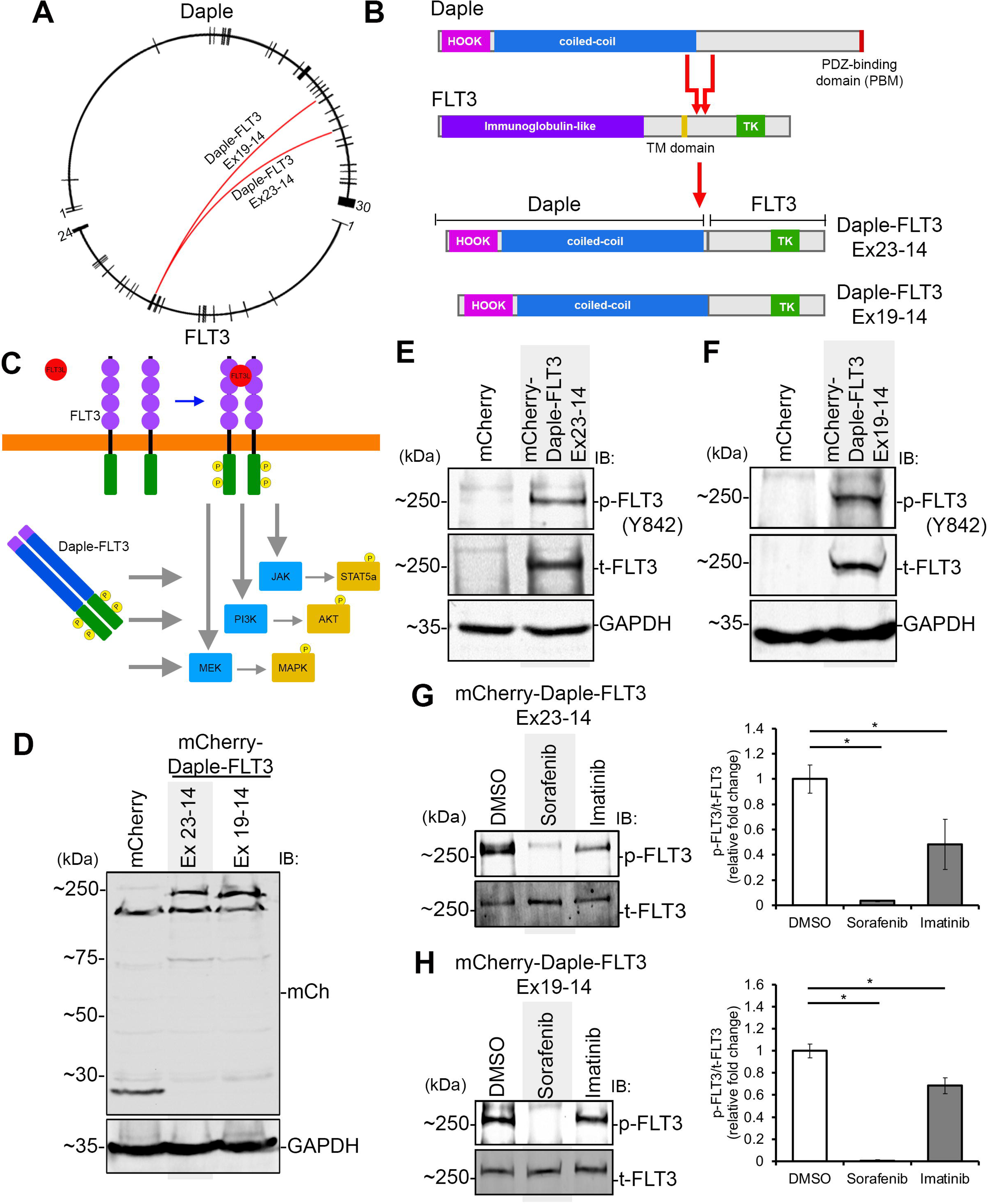
A constitutively active kinase domain is present on Daple-FLT3. **A)** Circos plot showing Daple and FLT3 transcripts. Ticks represent exons and line in between represent intronic region. Rearrangement point is illustrated through connecting line for Daple-FLT3 Ex23-14 and Ex19-14 (*red*). **B)** Illustration of protein domains found on Daple, FLT3, and the two Daple-FLT3 fusion products. TK, tyrosine kinase. **C)** Schematic of FLT3 receptor activation by its ligand FLT3L and the corresponding downstream activity of JAK/STAT5a, PI3K/AKT, and MEK/MAPK. Daple-FLT3, without an extracellular ligand binding domain, still activates STAT5a, AKT, and MAPK through its active tyrosine kinase domain. **D)** Immunoblot for mCherry (mCh) in K562 cell lysates ectopically expressing mCherry or the mCherry-tagged Daple-FLT3 gene fusion. **E-F)** K562 cells expressing mCherry or mCherry-tagged Daple-FLT3 (Ex23-14, *E*, and Ex19-14, *F*) were harvested and cell lysates were probed for phosphorylated FLT3 (Y842) and total FLT3. GAPDH was used for loading control. **G-H)** K562 cells expressing mCherry-tagged Daple-FLT3 (Ex23-14, *G*, and Ex19-14, *H*) were treated with DMSO, Sorafenib, or Imatinib for 24hours. After treatment, cells were collected for cell lysates and analyzed by immunoblotting for phosphorylated FLT3 (Y842) and total FLT3. Bar graphs on right show the relative phosphorylated to total FLT3 levels compared to DMSO control. Quantification was performed using densitometry on three independent repeat and significance was determine using student’s t-test (*, p<.05).

Oncogenic FLT3 alterations typified by FLT3-ITD have been shown to possess an active kinase and trigger pathways including STAT5a, AKT, and MAPK^11^. These pathways are known to drive cancer progression, including hematologic malignancies^12–14^. Cells with Daple-FLT3 are responsive to the first-generation FLT3 inhibitor sorafenib; furthermore, when Daple-FLT3 was expressed in Ba/F3, the fusion oncogene was strong enough to confer IL-3 independence^9,10^. Transformation into this factor independent state is typically reliant on elevated STAT5a^15^. These observations with Daple-FLT3 suggest that the FLT3 kinase is active when partnered to Daple; however, there is a lack of direct evidence demonstrating kinase activation or the subsequent elevation in signaling pathways (Figure 1C).

Forced oligomerization mediated through interaction domains, such as coiled-coil domains, are often seen in gene fusions^16–19^. In such cases, two (or more) kinase domains are brought together via hydrophobic interaction between coiled-coil domains and oriented in a fashion favoring autophosphorylation and activation. Daple is a coiled-coil domain containing protein and a regulator of Wnt signaling through its c-terminal PDZ-binding motif (PBM) interacting with Dvl. Its coiled-coil domain is rich in leucine repeats and has been shown to oligomerize^20,21^. Therefore, it is feasible that this domain is responsible for oligomerizing Daple-FLT3 in cells and plays a role in the kinase activation.

Motivated to understand how the Daple-FLT3 oncoprotein functions in modulating cellular signaling, we expressed this fusion oncoprotein in K562 cells, a cell line that has been previously used to study FLT3 and FLT3-ITD signaling^22^, and show that the STAT5a, AKT, and MAPK signal transduction pathways are all activated. This signaling pathway elevation is due to an active kinase domain on the oncoprotein and is independent of ligand binding. Both the kinase and signaling pathway activation can be modulated by tyrosine kinase inhibitors, aligning with the favorable response patients have in response to sorafenib. Furthermore, cells expressing the Daple-FLT3 fusions also showed response to the more specific second-generation FLT3 inhibitor quizartinib. Finally, we show that Daple contributes to the oncoprotein by enhancing kinase activation (through forced oligomerization) and localizing the protein to the pericentriolar space, although its removal could not completely abolish kinase activity.

## Methods

### Cell lines, culture methods, plasmid incorporation, and TKI treatment

K562 and Jurkat cells were cultured using RPMI media containing 10% FBS. To deliver plasmids into K562 cell, electroporation was performed using a Mirus EZPorator. Treatment with tyrosine kinase inhibitors was performed 24 hours post electroporation by replacing media with fresh media containing pharmacological inhibitors (10 μM drug concentration). Cells were treated for 24 hours (48 hours post electroporation) before harvesting for cell lysates. HEK293T cells were cultured using DMEM media containing 10% FBS and plasmids were delivered into cells using PEI transfection.

### Plasmid constructs

Cloning of Daple-FLT3 construct was carried out using Gateway cloning. Daple-FLT3 was PCR amplified, gel extracted and then used in a BP Clonase II (Thermo Fisher Scientific) reaction with pDONR221. The generated entry vector was then used in a LR Clonase II (Thermo Fisher Scientific) reaction with pDEST-CMV-N-EGFP^23^, pDEST-CMV-N-mCherry^23^, or pGCS-N1(6xMYC)^24^ to generate the plasmids for ectopic expression.

### Cell imaging and immunostaining

Following electroporation, cells were attached to poly-L-lysine coated coverslips for 60mins. For live-cell imaging, cell culture media was replaced with media containing Hoechst 33342 to stain nuclei and then immediately imaged on an APEXVIEW APX100 Benchtop Fluorescence Microscope (Evident Scientific). For immunostaining experiments, attached cells were fixed with 4% paraformaldehyde for 20mins, washed with 1XPBS, and then blocked with BSA (2mg/ml in PBS). For permeabilization, Triton Tx-100 was added to the blocking solution. Antibodies were diluted into blocking solution and incubated onto cells for 60mins. This was followed by washing off the excess antibody and then incubating with secondary antibody solution containing DAPI. Secondary antibody was then washed off and coverslip was mounted onto ProLong Gold antifade reagent (Thermo Fisher Scientific) before imaging.

### Recombinant Protein Purification

GST and HIS-tagged proteins were expressed in E. coli stain BL21 (DE3) and cultured for protein purification. Cultures were induced using 0.5mM IPTG overnight at 18°C. Cells were then pelleted and resuspended in GST lysis buffer (25 mM Tris-HCl, 150 mM NaCl, 1 mM EDTA, 20% (vol/vol) glycerol, 1% (vol/vol) Triton X-100, 150uM PMSF, 0.2mg/ml lysozyme, 1ug/ml DNase, and 1mM DTT, pH 7.5) or HIS lysis buffer (50mM Tris, 300mM NaCl, 10mM Imidazole, 1% (vol/vol) Triton X-100, 150uM PMSF, 0.2mg/ml lysozyme, 1ug/ml DNase, and 1mM DTT, pH 7.5). Afterwards, cells were then lysed by sonication, and lysates were cleared by centrifugation at 10,000 X g at 4°C for 20 mins. Proteins in supernatant were then affinity purified using Pierce GST Agarose (or Ni-NTA Resin for HIS-tagged purification), followed by elution, and overnight dialysis in TBS.

### In Vitro GST-Pulldown and Co-immunoprecipitation (CoIP) Assays

Purified GST-tagged proteins were immobilized onto GST beads and incubated with binding buffer (50 mM Tris-HCl (pH 7.4), 100 mM NaCl, 0.4% (v:v) Nonidet P-40, 10 mM MgCl_2_, 5 mM EDTA, 2 mM DTT) for 60 mins at 4°C. After binding, protein bound beads were washed, and the GST-protein bead complex was then incubated with HIS-tagged protein for 3 hours at 4°C. After binding, bound complexes were washed followed by elution through boiling at 100°C in sample buffer for SDS-PAGE analysis.

For CoIP assays, cells were lysed in mammalian cell lysis buffer (20 mM HEPES, 5 mM Mg-acetate, 125 mM K-acetate, 0.5% Triton X-100, 1mM DTT, 2X protease and phosphatase inhibitor, pH=7.4), clarified using centrifugation, and then cleared lysates were incubated with capture antibodies for 3 hours at 4°C. Afterwards, Protein A beads were added to capture the antibody bound protein complexes. Bound proteins were then eluted through boiling at 100°C in sample buffer for SDS-PAGE analysis.

### Proximity Ligation Assay (PLA)

PLA was performed using the Duolink® In Situ Proximity Ligation Assay Red Kit (Sigma-Aldrich), following the manufacturer’s protocol with minor modifications. Briefly, cells were cultured on poly-L-lysine-coated glass coverslips, fixed with 4% paraformaldehyde for 10 minutes at room temperature, and then permeabilized with 0.1% Triton X-100 in PBS for 10 minutes. Following permeabilization, coverslips were incubated in Duolink® blocking solution for 1 hour at 37 °C in a humidity chamber. Cells were then incubated with primary antibodies diluted in Duolink® antibody diluent, incubated for 90 minutes at room temperature, washed, and then incubated with the anti-mouse PLUS and anti-rabbit MINUS secondary antibodies diluted in antibody diluent for 1 hour at 37 °C. After incubation, coverslips were washed and then proceeded into a ligation reaction for 30 minutes at 37 °C. Myc antibody was purchased from Proteintech (16286-1-AP) and EGFP antibody was purchased from ThermoFisher (MA515256)

Following ligation, coverslips were washed and incubated with the amplification solution for 100 minutes at 37 °C in the dark. After amplification, samples were washed twice, mounted onto microscope slides using Duolink® In Situ Mounting Medium with DAPI, and sealed with nail polish. PLA signals (red fluorescent puncta indicating protein–protein proximity) and nuclear staining (DAPI) were visualized by fluorescence microscopy using APEXVIEW APX100 Benchtop Fluorescence Microscope.

## Results

### Expression of Daple-FLT3 into K562 cells shows kinase activation and activates STAT5, AKT, and MAPK pathways

To determine whether the chimeric Daple-FLT3 protein contains an active kinase, we first electroporated plasmids encoding the Daple-FLT3 fusion proteins (tagged to mCherry) into the myelogenous leukemia cell line K562, followed by immunoblotting of the cell lysates to confirmed expression and activation. Through immunoblotting for the mCherry tag, we see that these cells express the full-length protein (Figure 1D). Using the same lysates, we probed for phosphorylation using a phosphorylation site specific antibody to FLT3 (Y842). This key phosphorylation site is commonly used to indicate kinase activation^25^. In these lysates, we successfully detected phosphorylation for the Daple-FLT3 fusion proteins (Figure 1E and F).

The first-generation FLT3 inhibitor sorafenib have been shown to create a favorable response in a JMML patient with Daple-FLT3^9^. Therefore, we wanted to see what happens when K562 cells expressing Daple-FLT3 are treated with sorafenib. We also included imatinib in our studies because this inhibitor was shown to have favorable response in patients with Daple-PDGFRB fusion^26–28^. When cells expressing Daple-FLT3 were treated with sorafenib or imatinib, phosphorylation of Daple-FLT3 was decreased, with sorafenib being more efficient in decreasing phosphorylation compared to imatinib (Figure 1G and H).

Next, we set out to investigate if STAT5a, AKT, and MAPK signals are also activated in cells expressing the Daple-FLT3 protein. Cell lysates from K562 cells overexpressing the fusion oncoprotein were analyzed for phosphorylated STAT5a, AKT, and MAPK by western blotting as done above (Figure 2A and B). All three signaling pathways are significantly upregulated when compared to cells only expressing the mCherry tag control. Similarly, to further confirm that activation of the signaling pathways was dependent on tyrosine kinase activity, we asked if the tyrosine kinase inhibitors sorafenib or imatinib can prevent FLT3 activation and decrease phosphorylated STAT5a, AKT, and MAPK levels in these cells. We observed sorafenib decreasing phosphorylated STAT5a and AKT levels in both set of cells carrying Daple-FLT3. MAPK also showed decreased levels, but only with one of the Daple-FLT3 mutants (Figure 2C and D). With imatinib treatment, there was a decrease in STAT5a phosphorylation in Daple-FLT3 Ex23-14, but not Ex19-14. No statistically significant changes in phosphorylated AKT or MAPK was observed, but interestingly, these levels appear to increase (Figure 2C and D).

**Figure 2.**
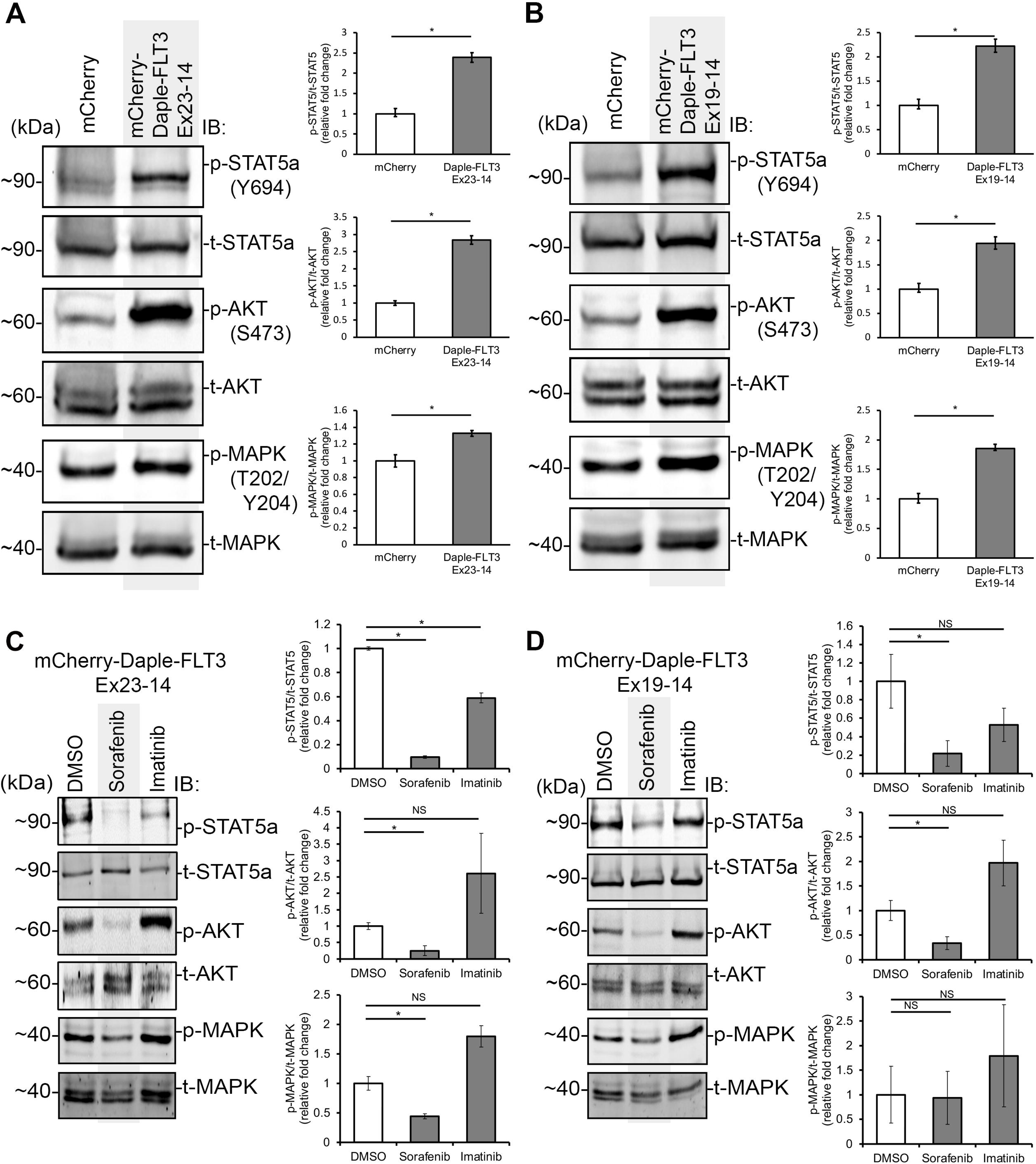
Daple-FLT3 elevates signaling pathways and is blocked by tyrosine kinase inhibitors. **A-B)** K562 cells expressing mCherry or mCherry-tagged Daple-FLT3 (Ex23-14, *A*, and Ex19-14, *B*) were harvested and cell lysates were probed for phosphorylated STAT5a (Y694), AKT (S473), and MAPK (T202, Y204) as well as the respective total protein. Bar graphs on right show the relative phosphorylated to total protein levels compared to mCherry control. Quantification was performed using densitometry on three independent repeat and significance was determine using student’s t-test (*, p<.05). **C-D)** K562 cells expressing mCherry-tagged Daple-FLT3 (Ex23-14, *C*, and Ex19-14, *D*) were treated and collected as above. Cell lysates were analyzed by immunoblotting for phosphorylated STAT5a (Y694), AKT (S473), and MAPK (T202, Y204) as well as for the respective total protein. Bar graphs on right show the relative phosphorylated to total protein levels compared to DMSO control. Quantification was performed using densitometry on three independent repeat and significance was determine using student’s t-test (*, p<.05. NS, not significant).

Second-generation FLT3 inhibitors such as quizartinib have more specificity and potency than first-generation inhibitors (i.e. sorafenib). Therefore, we wanted to test the response of cells ectopically expressing either of the Daple-FLT3 mutant proteins to quizartinib. Similar to sorafenib treatment, quizartinib also showed decrease phosphorylated STAT5a, AKT, and MAPK levels (Figure 3A and B).

**Figure 3.**
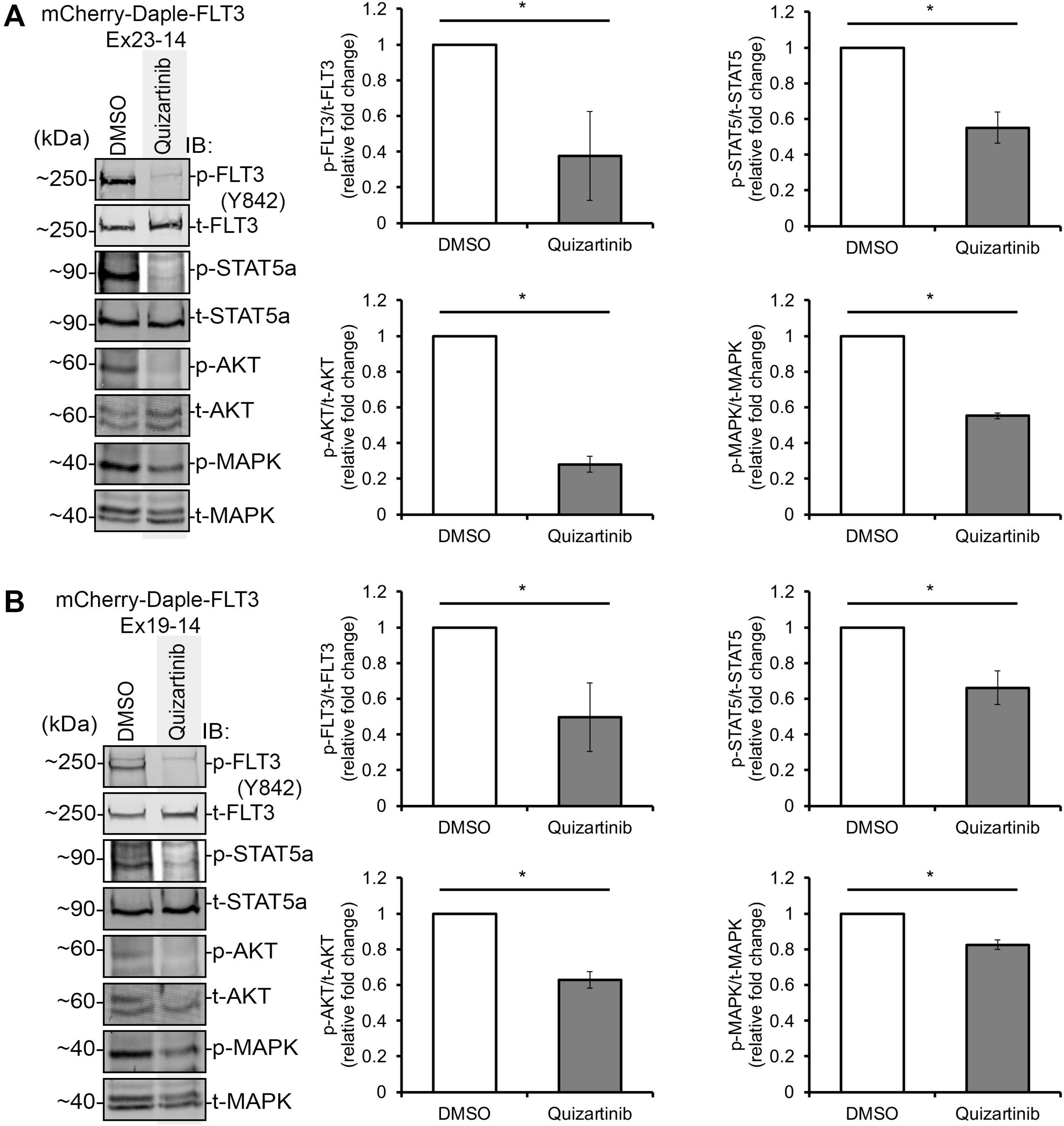
Targeting Daple-FLT3 using the FLT3 specific inhibitor quizartinib. **A and B)** K562 cells expressing mCherry-tagged Daple-FLT3 (Ex23-14, *A*, and Ex19-14, *B*) were treated with the FLT3 specific inhibitor quizartinib and collected for SDS-PAGE. Cell lysates were analyzed by immunoblotting for phosphorylated FLT3 (Y842), STAT5a (Y694), AKT (S473), and MAPK (T202, Y204) as well as for the respective total protein. Bar graphs on right show the relative phosphorylated to total protein levels compared to DMSO control. Quantification was performed using densitometry on three independent repeat and significance was determine using student’s t-test (*, p<.05).

### Daple-FLT3 shows subcellular localization

Chromosomal rearrangement leading to protein domain loss can affect its spatial and temporal regulation^29,30^. The Daple-FLT3 fusion protein is lacking the C-terminal portion of Daple, which contains its heterotrimeric G-protein binding region^21^ and PDZ-binding motif^20^ (Figure 1B). The FLT3 portion on Daple-FLT3 lacks the ligand binding domain and the transmembrane domain. To investigate the consequences of such losses on Daple-FLT3, we expressed mCherry tagged Daple-FLT3 in cells and observed for localization under live fluorescent microscopy. We see that the tagged fusion proteins exhibit a cytosolic and subcellular localization pattern, whereas the mCherry only tag control revealed a cytosolic localization pattern (Figure 4A and Supplemental Figure 1A).

**Figure 4.**
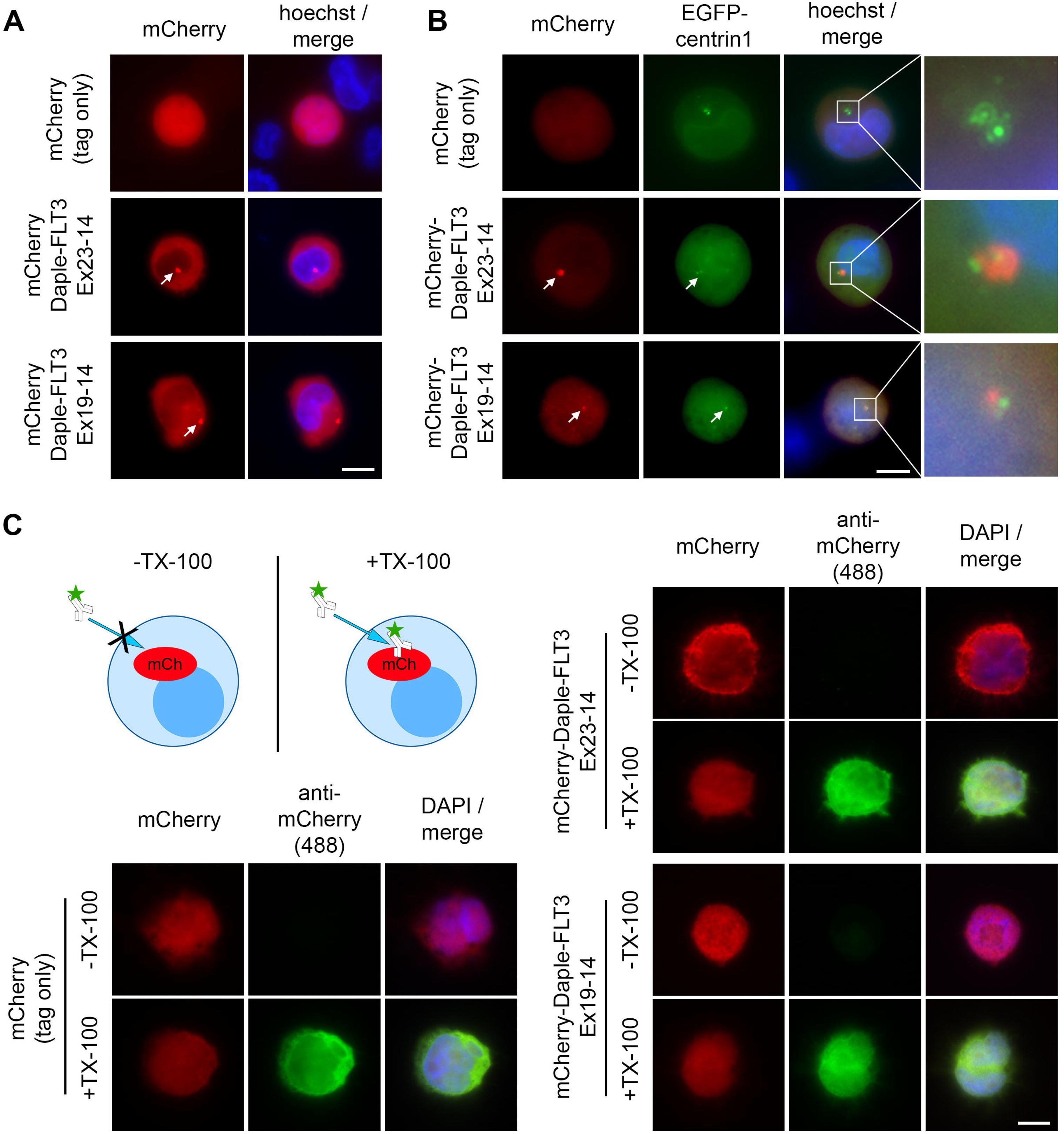
Daple-FLT3 localizes to the centrosome. **A)** Live-cell imaging of K562 cell expressing mCherry or the mCherry-Daple-FLT3 constructs. Nuclei were imaged with Hoechst staining. Scale bar, 10μm. **B)** EGFP-centrin1 was co-expressed with either mCherry-Daple-FLT3 Ex23-14 or mCherry-Daple-FLT3 Ex19-14 and imaged live. Nuclei were imaged with Hoechst staining. Scale bar, 10μm. **C)** Illustration depicts anti-mCherry only being able to enter cells and bind mCherry epitope when cells are permeabilized with TX-100 after fixation with paraformaldehyde. Fixed cells were immunostained using an anti-mCherry antibody, followed by binding to an AlexaFluor-488 conjugated secondary antibody. Natural fluorescence of mCherry is preserved after fixation. Nuclei were imaged with DAPI staining. Scale bar, 10μm.

Prior work has shown that Daple can subcellularly localize to the pericentriolar region in various epithelial cell lines^31–33^. Furthermore, the coiled-coil domain alone is sufficient in localizing to this region^33^. Because the Daple-FLT3 gene fusions retain the Daple coiled-coil domain, we asked if this chimeric product also localizes to the pericentriolar space. To address this, cells expressing the tagged oncofusion protein were co-expressed with EGFP tagged Centrin-1 to mark the centriolar region. As expected, we see that both Daple-FLT3 localizes along with EGFP-Centrin-1 (Figure 4B). Staining for the centrosomal marker γ-tubulin further confirms the localization of the oncofusion protein to the pericentriolar space (Supplemental Figure 1B).

Next, because the Daple-FLT3 protein is lacking the FLT3 transmembrane domain and varies in localization from FLT3 and FLT3-ITD (Supplemental Figure 2), we wanted to confirm that Daple-FLT3 is intracellularly localized and absent from the plasma membrane. To this end, we electroporated the mCherry tagged gene fusion into cells and performed immunostaining using a mCherry antibody in either TX-100 permeabilized or unpermeablized cells. Positive immunostaining for the mCherry antibody (revealed though a AlexaFluor-488 conjugated secondary antibody) was only detected in the TX-100 permeabilized condition (Figure 4C). Because PFA fixation preserved the natural fluorescence of the mCherry molecule, we were able to identify cells that were successfully electroporated. This intracellular localization of Daple-FLT3, taken together with kinase activation without ligand treatment, suggest that Daple-FLT3 is constitutively activated inside the cell.

### Daple-FLT3 oligomerizes in cells

Coiled-coil domains are frequently involved in protein-protein interactions—including oligomerization with other coiled-coils^17,34^. When rearranged to kinases, this domain can facilitate transphosphorylation by self-binding into dimers or larger oligomers^17^. Daple’s coiled-coil domain is rich in leucine residues and ordered in a heptad repeat. Studies using the mouse Daple protein demonstrated that a leucine rich stretch of the coiled-coil domain partakes in protein oligomerization^20^. Protein alignment between Daple in mouse and human reveal a high conservation of the leucine residues across this leucine rich region—with human Daple containing one less leucine within the heptad repeats (Supplemental Figure 3). This leucine rich stretch of Daple is also present in the Daple-FLT3 oncoproteins. Therefore, we wanted to test the possibility that Daple-FLT3 can oligomerize.

Because prior oligomerization studies were performed with mouse Daple, we first set out to test if the human Daple can oligomerize^20^. To this end, we utilized a GST pulldown assay between GST-tagged and HIS-tagged Daple coiled-coil domain. When this was carried out, we observed that GST-tagged Daple coiled-coil domain was indeed able to pulldown HIS-tagged Daple coiled-coil domain out of solution (Figure 5A). This confirms that a direct interaction occurs between the two fragments.

**Figure 5.**
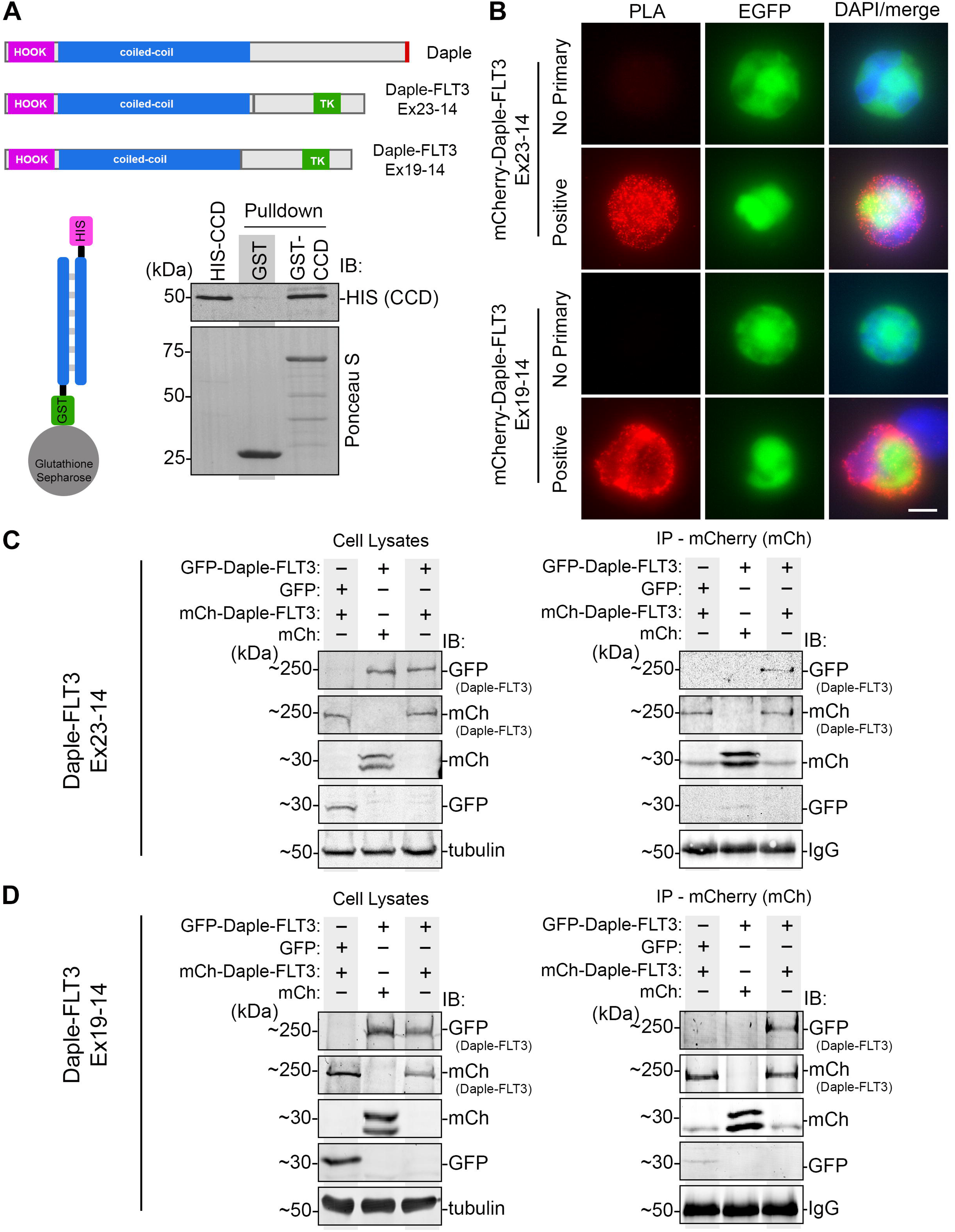
Daple-FLT3 oligomerizes in cells. **A)** Pull-down assays using purified GST-tagged Daple coiled-coil domains (CCD) immobilized on glutathione beads and soluble recombinant His-Daple-CCD. Bound Daple-CCD was determined by immunoblotting. **B)** Proximity-Ligation Assay in K562 cells expressing myc-Daple-FLT3 and EGFP-Daple-FLT3. Scale bar, 10μm. **C-D)** Co-immunoprecipitation assays investigating the binding between mCherry-Daple-FLT3 and EGFP-Daple-FLT3 (Ex23-14, *C*, and Ex19-14, *D*). Proteins were exogenously expressed in HEK293T cells followed by cell lysis and IP using anti-mCherry antibody.

Next, we wanted to determine if Daple-FLT3 oligomerizes in cells. To test this possibility, we performed a proximity ligation assay (PLA) on K562 cells co-expressing myc-tagged and EGFP-tagged Daple-FLT3 (Figure 5B). We observed positive punctate signals throughout the cell, suggesting that oligomerization occurs and that this was ubiquitous throughout the cell. Coimmunoprecipitation experiments confirmed such findings (Figure 5C and D). In these experiments, both mCherry-tagged and EGFP-tagged Daple-FLT3 were co-transfected into HEK293 cells and lysates were incubated with a mCherry antibody. Immunocomplexes were then analyzed by SDS-PAGE and immunoblotting. We observed that EGFP-Daple-FLT3 (but not EGFP alone) complexes with mCherry-tagged Daple-FLT3. The mCherry tag alone neither precipitated with EGFP or EGFP-Daple-FLT3 ruling out any non-specific interactions between tags. Collectively, these findings demonstrate that Daple-FLT3 indeed oligomerize in cells.

### Daple Coiled-coil domain maximizes kinase activation and is necessary for localization

The coiled-coil domain on Daple is sufficient for subcellular localization to the pericentrosomal space and oligomerization^35^. To determine the function of Daple on the Daple-FLT3 fusion, we deleted the Daple portion and ectopically expressed the FLT3 region into K562 cells (Figure 6A). Using live fluorescent microscopy, we saw that the kinase domain was only observed within the cytoplasm. This contrasted with the full-length fusion protein, which showed both cytoplasmic and subcellular localization (Figure 6B). Next, we asked if Daple contributed to kinase activation by assaying for phosphorylation of the kinase domain when expressed in K562 cells. Without the Daple partner gene, FLT3 phosphorylation was significantly decreased compared to full-length Daple-FLT3 (Figure 6C and D). It is worth noting that FLT3 phosphorylation, albeit weak, could still be detected. We confirmed this through immunoprecipitation experiments to enrich for the ectopically expressed protein (Supplemental Figure 4). Coinciding with less FLT3 phosphorylation, STAT5a and AKT activation were also significantly decreased. MAPK, however, did not show significant changes (Figure 6E and F). Taken together, these results suggest that Daple is not only essential for maximal activation of the tyrosine kinase, but also necessary for localization of the protein.

**Figure 6.**
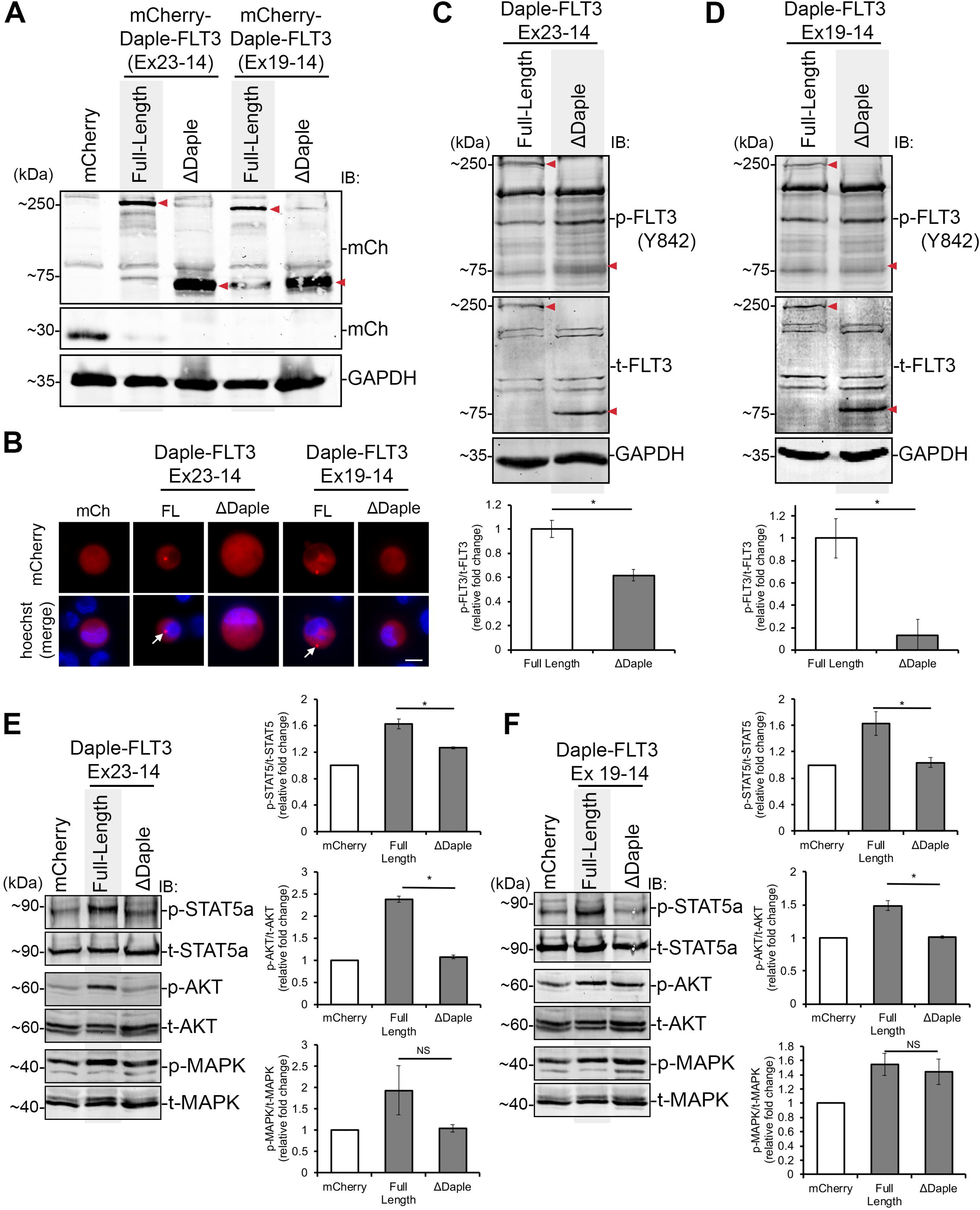
Maximal kinase activation and signaling is dependent on the Daple partner gene. **A)** Immunoblot for mCherry (mCh) in K562 cell lysates ectopically expressing mCherry, mCherry-tagged Daple-FLT3, or the Daple deleted mutant (ΔDaple). **B)** Live-cell imaging of K562 cell expressing mCherry, both mCherry-Daple-FLT3 constructs, as well as their respective Daple deleted (ΔDaple) fusion protein. Scale bar, 10μm. Side panels (*right*) show digitally zoomed region. **C-D)** K562 cells expressing mCherry, the mCherry-tagged Daple-FLT3 (Ex23-14, *C*, and Ex19-14, *D*), or their respective Daple deleted mutant (ΔDaple) were harvested, lysed, and cell lysates were immunoblotted for phosphorylated FLT3 (Y842) and total FLT3. GAPDH was used for loading control. Arrows point to FLT3 band of interest. **E-F)** K562 cells expressing mCherry, mCherry-tagged Daple-FLT3 (Ex23-14, *E*, and Ex19-14, *F*), as well as their respective Daple deleted (ΔDaple) fusion protein was harvested, lysed, followed by immunoblotting the lysates for phosphorylated STAT5a (Y694), AKT (S473), and MAPK (T202, Y204) as well as the respective total protein. Bar graphs on right show the relative phosphorylated to total protein levels normalized to mCherry control. Quantification was performed using densitometry on three independent repeat and significance was determine using student’s t-test (*, p<.05, NS, not significant).

## Discussion

Gene fusions play an integral role in the development of many cancers. It is estimated that 20% of cancers contain some form of gene fusion, with the rate being higher (∼30%) in hematological malignancies^36,37^. Here, we demonstrate that Daple-FLT3, a rare but recurrent gene fusion in hematological malignancies, carries a constitutively active kinase that activates downstream pathways including STAT5a, AKT, and MAPK. Moreover, the kinase and signaling pathways can be targeted using tyrosine kinase inhibitors such as sorafenib and quizartinib. Interestingly, we observe that imatinib had less effect on kinase activation compared to sorafenib. Although the reason for this observation with imatinib remains unknown, this does align with prior observation that FLT3 kinase activity can render K562 cells insensitive to imatinib. Furthermore, clinical evidence show that imatinib is generally ineffective in FLT3-altered cancers^38^.

Kinase fusion to an oligomerization domain, such as coiled-coil domains, is found in over 60% of gene fusions. This is significant when one considers that this domain only makes up 9% of the human proteome^39^. The function of these fusion oncoproteins typically leads to enforced oligomerization and kinase activation^17^, although this is not always necessary for activation^40^. Interestingly, while the Daple gene fusion does indeed oligomerize, removing the Daple portion and solely expressing the FLT3 portion, does not completely abolish kinase activation or signaling. Thus, although Daple is dispensable for the kinase activation, it is required for maximal activation.

The observation that expression of just the FLT3 kinase domain is sufficient for kinase activation is interesting. This leads to the possibility that chromosomal rearrangement between Daple and FLT3 can lead to hematological defects through two mechanisms: 1) through promoter hijacking, where the FLT3 kinase domain is driven to a level higher than its own native promoter, and 2) where the coiled-coil domain promotes dimerization and enhances activation beyond what is achieved through stochiometric interaction alone. These two mechanisms are not mutually exclusive; therefore, it is possible that each play a contribution. Further work will be necessary to tease out the individual contributions.

In addition to enhancing kinase activation, Daple’s coiled-coil domain also facilitates the localization of the gene fusion to the pericentriolar space. Access to its substrate determines kinase signaling activity, therefore, its subcellular localization can greatly influence its outcome^41–43^. FLT3 is a receptor tyrosine kinase normally found on the cell surface; however, FLT3-ITD have been found to be internally localized to Golgi and endoplasmic reticulum (ER)^44^. FLT3-ITD localization to each of these compartment influences it’s signaling outcome, with STAT5a activation coming from the ER, while AKT and MAPK arising from the Golgi. Unlike FLT3-ITD, Daple-FLT3 lacks a transmembrane domain, and is localized to the cytosol and pericentriolar space. This observation leads to the interesting notion that STAT5a, AKT, and MAPK may be activated through a distinct mechanism from other FLT3 mutations; however, this remains to be tested. Moving forward, it will be informative to dissect the similarities and differences between membrane bound FLT3 mutants in leukemia to the Daple-FLT3 fusions. Understanding such differences may unveil areas for personal therapy.

Secondary mutations can occur in the kinase domain, rendering it resistant to TKI therapy. In such cases, switching to a different TKI, increasing dosage, or using a combinatory strategy with multiple TKI is performed^45,46^. However, off-target effects and the patient’s overall health can limit treatment options. Because Daple is necessary for maximal activation of the kinase domain, it may be possible to target Daple’s coiled-coil as an alternative or complementary treatment route. Such strategies have been proposed with other gene fusions^47^.

During chromosomal rearrangements, one copy of a gene is lost. In the case of Daple rearrangement to FLT3, this would mean one copy of the normal Daple gene is lost. Daple is expressed during interphase and is an important regulator of Wnt signaling and primary cilia orientation^21,48,49^. How loss of Daple will impact Wnt signaling in hematopoietic cell is an avenue to be explored. Furthermore, both the Wnt and Hedgehog signaling pathways are highly regulated by the primary cilia, and they play an important role in both normal and malignant cells^50,51^. How the constitutive activation of the FLT3 kinase at the centrioles, due to the Daple-FLT3 fusion, affect these pathways remain an area to be studied.

This work demonstrates that chromosomal rearrangement between Daple and FLT3 can lead to a stable product with an elevated kinase activity. Furthermore, modulating Daple can influence such activity. Chromosomal rearrangement between Daple to other kinases have been identified, such as to PDGFRB and JAK2^26–28,52^. These rearrangements also carry the same domain structure in that Daple’s HOOK and coiled-coil domain (albeit to varying lengths) is fused to the entire kinase domain. Whether such product can lead to the same localization pattern and signaling activity as Daple-FLT3 remains to be tested. If so, this creates the possibility of targeting Daple as a common therapeutic route and, thus, warrant further investigation.

## Acknowledgments

This work was supported by NIH grant R16GM146720 and a CSUPERB New Investigator grant to J.E. A.I was supported by a NIH training grant T32GM137812 during the course of this work. The content is solely the responsibility of the authors and does not necessarily represent the official views of the National Institutes of Health. pEGFP-centrin-1 was a gift from Michel Bornens (Addgene plasmid # 72641). pDEST-CMV-N-EGFP and pDEST-CMV-N-mCherry were gifts from Robin Ketteler (Addgene plasmid #122842 and 123215). pGCS-N1(6xMYC) was a gift from Hai-Ning Du (Addgene plasmid # 85718).

## Authorship Contributions

J.E., D.K., and M.A. designed the studies. J.E., D.K., M.A., A.I., E.N., H.M., and K.A. performed experiments. E.S. generated and provided the plasmid used in studies. J.E. and D.K. wrote and revised the manuscript.

## Disclosure of Conflicts of Interest

The authors declare no competing financial interests.

## Notes

### Competing Interest Statement

The authors have declared no competing interest.

### Summary of Updates

Figure 3 revised and updated with new drug treatment. Supplemental files updated.

