## Supplemental Data for "Daple-FLT3 (CCDC88C-FLT3) gene fusion requires the coiled-coil domain for maximal activation and pericentrosomal localization"

#### Supplemental Figure 1

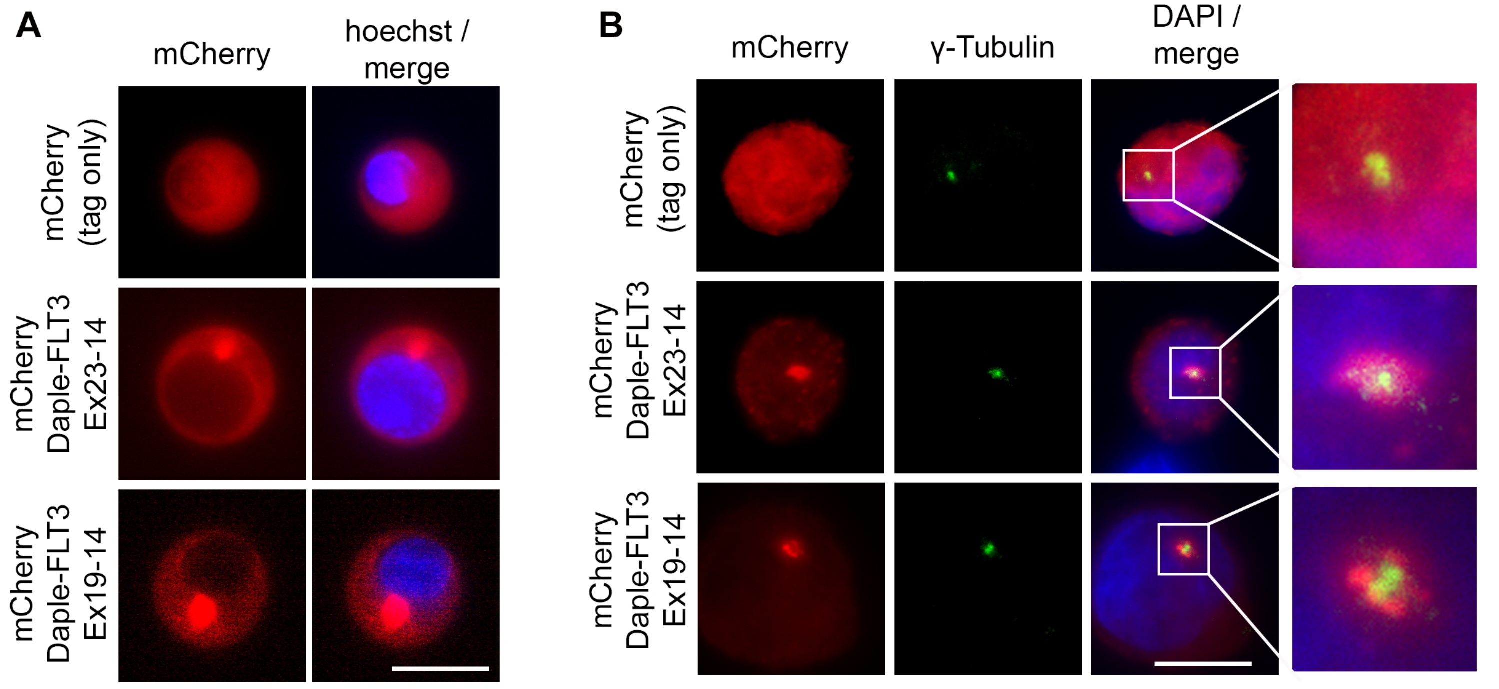

#### Supplemental Figure 2

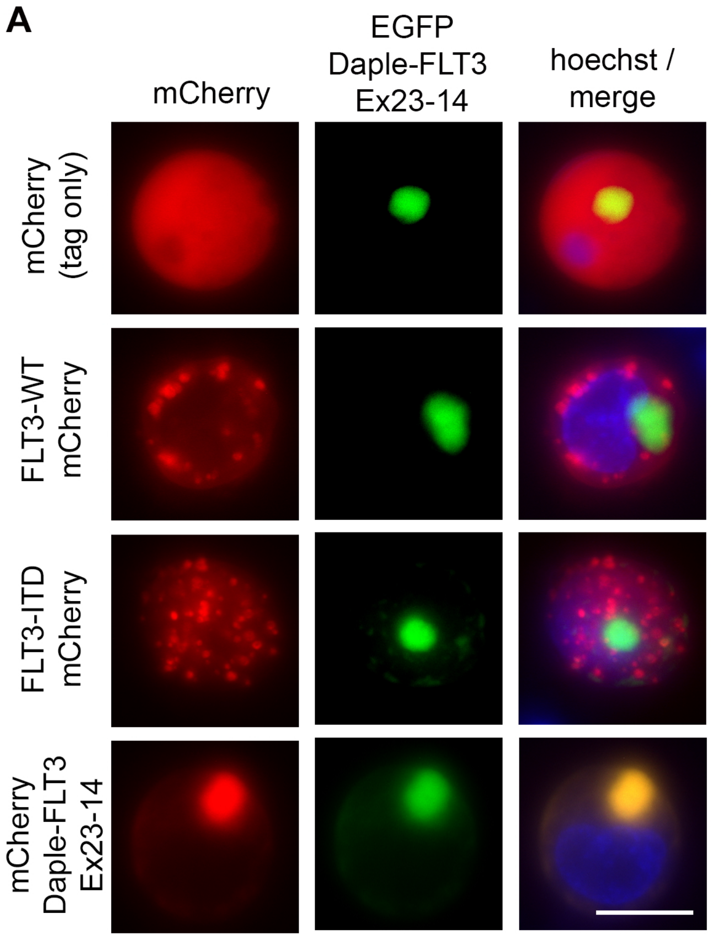


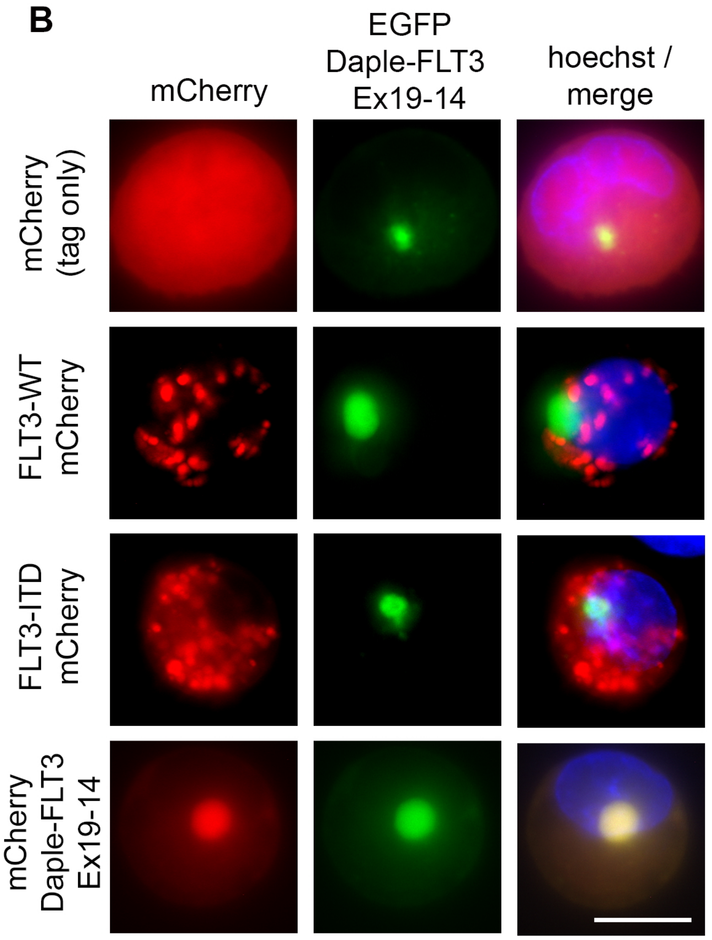

### Supplemental Figure 3

```
        abcdefg
mouse   LERSNAALQAERQLLKEQLQHLETQNVSFSSQILTLQKQSAFLQEHTTT-1101
human   LERNNAALQAEKQLLKEQLQHLETQNVTFSSQILTLQKQSAFLQEHNTT-1110
        ***.*******:***************:*****************.***

mouse   LQTQTAKLQVENSTLSSQNAALSAQYTVLQSQQAAKEAEHEGLQQQQEQ-1150
human   LQTQTAKLQVENSTLSSQSAALTAQYTLLQNHHTAKETENESLQRQQEQ-1159
        ******************.***:****:**.:::***:*:*.**:****

mouse   LAAVYEALLQDHKHLGTLYECQSSEYEALIRQHSCLKTLHRNLELEHKE-1199
human   LTAAYEALLQDHEHLGTLHERQSAEYEALIRQHSCLKTLHRNLELEHKE-1208
        *:*.********:*****:* **:*************************

mouse   LGERHGDLQQRKAELEELEKVLSTEREAL-1228
human   LGERHGDMLKRKAELEEREKVLTTEREAL-1237
        *******: :******* ****:******
```

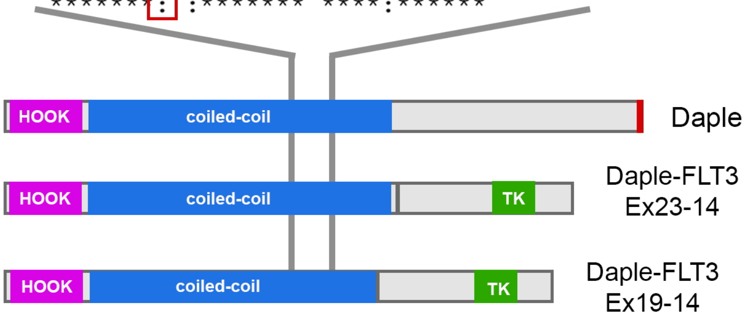

### Supplmental Figure 4

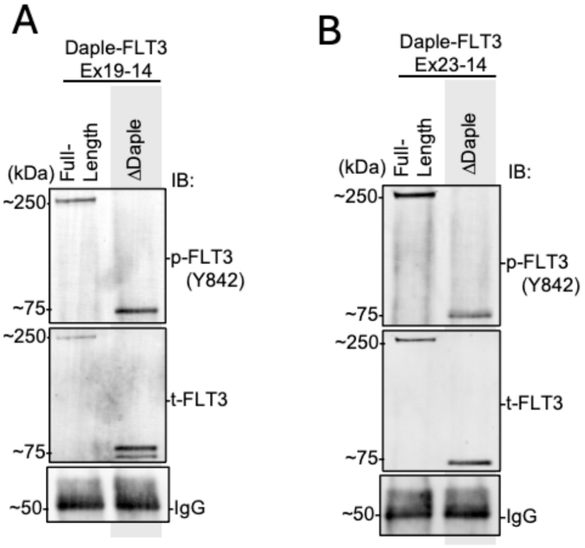

**Supplemental Figure 1. A)** Live-cell imaging of Jurkat cells expressing mCherry or the mCherry-Daple-FLT3 constructs. Nuclei were imaged with Hoechst staining. Scale bar, 10μm. **B)** Immunostaining using for mCherry (*red*) and γ-tubulin (*green)* in K562 cells expressing Daple-FLT3*.* Nuclei were imaged with DAPI staining. Scale bar, 10μm.

**Supplemental Figure 2. A and B)** Live-cell imaging of K562 cells co-expressing EGFP tagged Daple-FLT3 (Ex23-14, panel A, or Ex19-14, panel B) along with mCherry, mCherry tagged FLT3-wild type (WT), FLT3-ITD or the mCherry-Daple-FLT3 constructs. Nuclei were imaged with Hoechst staining. Scale bar, 10μm.

**Supplemental Figure 3.** *Top*, Mouse and human Daple amino acid sequence alignment on the leucine rich region. Leucine residues follow a heptad repeats (indicated *a* through *g*) with leucine, L, highlighted in the first position. Red box indicates where there is one less leucine residue on the human protein sequence. *Bottom*, the leucine rich region is also found on both Daple-FLT3 fusion oncoprotein.

**Supplemental Figure 4.** mCherry immunoprecipitation assays to enrich for mCherry-Daple-FLT3 full-length and Daple deleted mutant (Daple) (Ex19-14, *A*, and Ex23-14, *B*). After enrichment, eluted proteins were run on SDS-PAGE and immunoblotted for total and phosphorylated FLT3.
